# Contact-based kin discrimination is associated with specific surface lipids in the cannibalistic nematode *Pristionchus pacificus*

**DOI:** 10.64898/2026.01.13.699210

**Authors:** Fumie Hiramatsu, Desiree L. Goetting, Anna M. Kotowska, Nurit Zorn, Veeren M. Chauhan, James W. Lightfoot

## Abstract

Kin-recognition is widespread, yet its molecular basis remains poorly understood. In the predatory nematode *Pristionchus pacificus*, kin-recognition prevents cannibalism of close relatives and depends on the peptide SELF-1. Here, we show this behavior is robust to environmental stress but disrupted by a surfactant wash, implicating lipids or other amphiphilic molecules as necessary components for this mechanism. Using 3D-OrbiSIMS, we profiled the outer cuticle of kin-recognition defective *self-1* mutants and found these animals exhibit distinct surface lipids. Furthermore, analysis of surface chemistry defective *daf-22* mutants alongside *self-1;daf-22* double mutants revealed additional, non-overlapping surface lipid profiles, indicating that multiple pathways contribute to shaping surface lipid composition. Importantly, by combining automated behavioral tracking with state-based analysis, we show *daf-22* mutants are also kin-recognition defective. Together, these findings demonstrate that the composition of the nematode surface is required to maintain kin identity, with the cuticle acting as a signaling interface that regulates contact-dependent behaviors.

**Teaser (125 characters):** Surface lipid composition correlates with nematode kin-recognition signaling abilities.

## Introduction

The ability to identify and differentiate kin from unrelated individuals is deeply conserved across phylogeny. The perception of kin influences a wide variety of behaviors, ranging from boundary formation, aggregation, and cooperation in single-celled organisms (*1–3*), to mate preference, rearing of offspring, resource allocation, and even cannibalism in multicellular organisms (*4–7*). Kin recognition and its pervasive influence on behavior have long fascinated evolutionary biologists and neuroethologists, and yet, the molecular mechanism underlying the ability to discriminate kin has been challenging to decipher. Our clearest understanding of the molecular determinants of kin-determination mechanisms comes from studies on a select group of organisms. In yeast, kin discrimination is governed by the flocculation (FLO) genes and is polygenic and context dependent (*2*, *8*). In the social amoeba *Dictyostelium discoideum*, the gene pair TgrB1 and TgrC1 (Transmembrane glycoprotein involved in kin-recognition) encodes a highly polymorphic cell-surface receptor-like adhesion protein and its binding partner. Together with other Tgr family and accessory proteins, these loci enable these organisms to preferentially adhere to and cooperate with others expressing matching TgrB1/TgrC1 allele combinations (*9–11*). In marine invertebrates such as *Botryllus schlosseri*, *Hydractinia*, and sponges, colony fusion or rejection is regulated by the compatibility of gene products also encoded at highly polymorphic loci (*12–15*). Furthermore, in several multicellular species, secreted compounds, such as hydrocarbons in Drosophila (*16*), ants (*17*) and other insects (*18*) or odorants and pheromones in vertebrates (*19–21*), have emerged as key components of kin signaling, but the mechanism by which these molecules mediate kin recognition remains poorly understood.

Recently, nematodes have emerged as a powerful system for understanding the molecular mechanism underlying kin discrimination and its influence on social behavior. In particular, kin-recognition has been identified in several nematodes capable of predation. This includes many nematodes belonging to the Diplogastrid family such as species of *Pristionchus* (*22*, *23*) and the basal outgroup *Allodiplogaster sudhausi* (*24*), as well as taxonomically distinct predatory species including *Seinura caverna* (*25*). In these nematodes, robust kin signaling prevents their own progeny and closely-related conspecifics from being attacked and cannibalized. The most intensively studied of these nematode species is *Pristionchus pacificus,* where in addition to suppressing predatory feeding behavior, kin discrimination influences various other social interactions including aggregation and territorial behaviors (*26*, *27*). In *P. pacificus,* kin signaling requires the small peptide SELF-1, that is expressed in the hypodermal layer of the body wall and contains a hypervariable region necessary for its function (*22*, *23*). However, the role of the environmentally-exposed layers of the cuticle, and its vast array of surface-bound molecules, in establishing nematode identity remains unknown.

Here, we investigate the role of the cuticle and its outer surface for kin signaling. By exposing the cuticle to extreme external stressors, we show that kin signaling is robust to perturbation by nearly all surface treatments tested with the exception of a lipid-destabilizing surfactant. This suggests the presence of surface-associated lipids are necessary for maintaining this kin-determination system. Subsequently, the importance of the surface lipids is reinforced as kin-recognition defective *self-1* mutants also have an aberrant surface lipid profile. Furthermore, we investigate *daf-22* mutants that have a highly dysregulated surface chemistry (*28*), and find these are also kin-recognition defective. Therefore, mutants with an aberrant surface chemistry fail to generate an effective kin-signal, consistent with a role for the outer nematode surface as a signaling entity.

## Results

*P. pacificus* is an omnivorous nematode that supplements its bacterial diet with predatory feeding on the larvae of other nematodes (Fig. 1A). This predatory behavior is dependent on the presence of teeth-like-denticles (Fig. 1B) (*29*) that pierce through the outer cuticle of the prey and facilitate the consumption of the innards. Importantly, predation is not limited to just other nematode species as they also cannibalize con-specifics. However, a robust kin-recognition system protects close relatives from these predatory attacks (*22*, *23*). The propensity to cannibalize conspecifics can be quantified using previously established assays where adult predators are placed together with an abundance of juveniles in the absence of a bacterial food source. Subsequently, the number of corpses present are quantified as an indicator of successful predatory events (Fig. S1) (*30*, *31*). When predators are presented with genetically diverse conspecifics, abundant corpses can be observed while self-progeny are identified and spared from predation due to their robust kin-recognition system. Examples of the kin-recognition system can be observed in interactions between the main lab isolate PS312, used in most molecular studies, and the distantly related PS1843 (Fig. 1C, D) (*32*). As prey detection requires nose contact between the predator and potential prey (*33*, *34*), a kin-identity cue is likely generated through surface-bound signals associated with the *P. pacificus* cuticle. While it has been hypothesized that the presence of SELF-1 on the nematode surface may act as a kin signal, efforts to identify the peptide within the cuticle have so far been unsuccessful (*23*). Therefore, to investigate the role of the nematode surface in kin signaling, we attempted to disrupt any potential surface cues.

**Figure 1.**
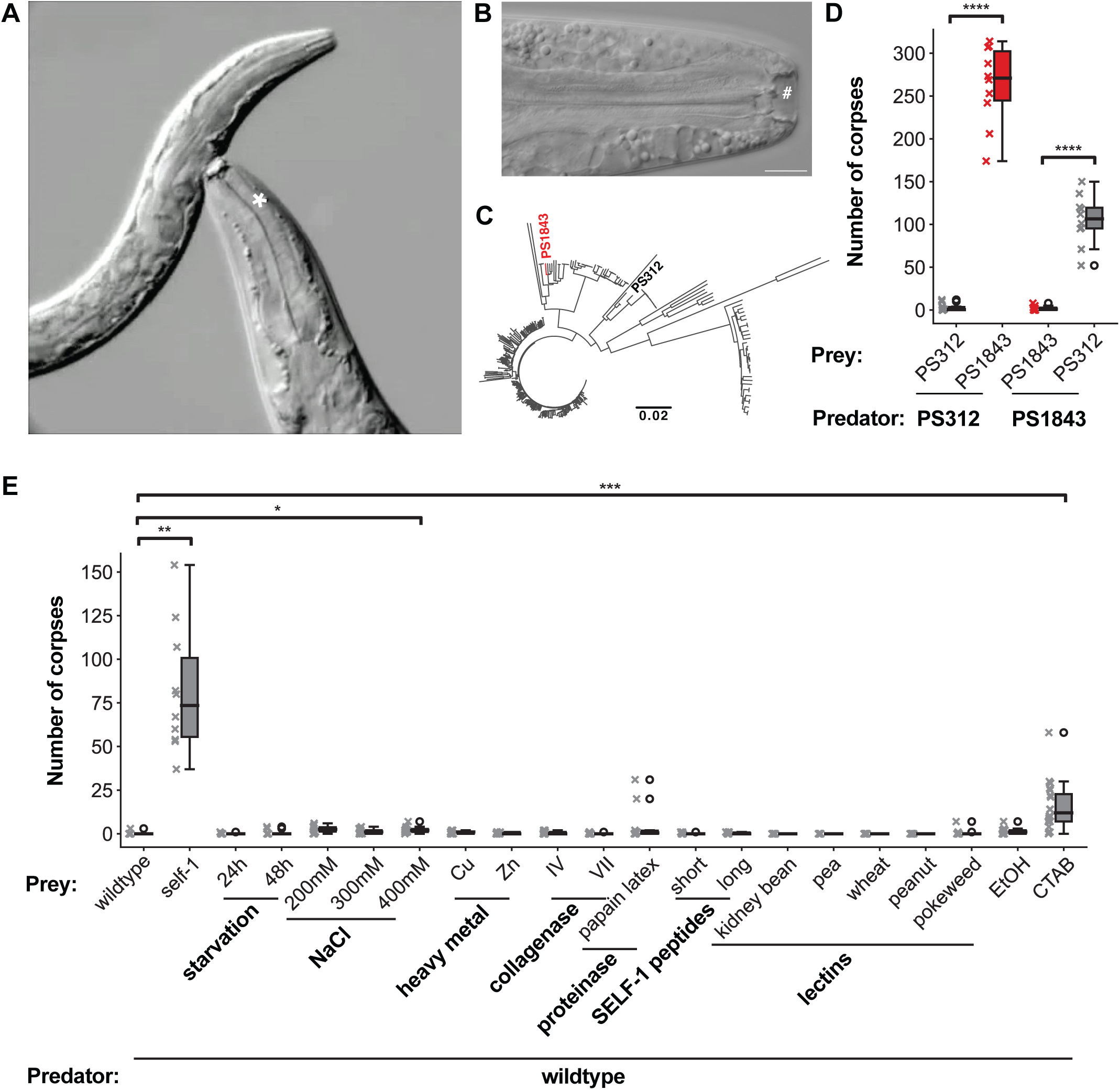
Robust kin-signaling prevents cannibalism. (A) An adult *P. pacificus* (*) hermaphrodite killing a *C. elegans* larvae. (B) DIC of *P. pacificus* mouth showing two teeth-like denticles (#) used for killing other nematodes. Scale bar = 10 µm. (C) Phylogenetic tree of *P. pacificus* representing the genetic relationship between 323 wild isolates. Two distantly related strains, PS312 and PS1843 are highlighted. Image adapted from Rödelsperger et al.,(*32*) . (D) Corpse assays assessing predatory interactions between adult predators and larval prey. PS312 and PS1843 are used as either predator or prey in reciprocal assays. Strains cannibalize one another but do not kill their own progeny. (E) Corpse assays of PS312 larvae treated with environmental stressors, then assessed for kin-recognition defects by challenging with PS312 predators. *self-1* mutant prey used as a control. For all corpse assays, each data point indicates the number of corpses present after 24 h in an individual trial of five predators on an assay plate saturated with juveniles. N=10 per condition, *p<0.05, **p<0.01, ***p<0.001 Mann-Whitney-U-Test with Bonferroni correction.

We first explored the robustness of the kin signal by challenging larvae from the wildtype strain PS312 with nutritional or environmental stressors. These included extremes of starvation, osmotic stress, and exposure to heavy metals known to disrupt the cuticle in several nematode species (*35*). We observed a very mild effect under high osmotic stress while kin recognition was robustly maintained under all other stresses (Fig. 1E). As SELF-1 is required for the kin signal (*23*), and we previously predicted that it may be present on the nematode surface, we investigated if the presence of additional SELF-1 peptide variants from other strains would disrupt the kin-signal. We therefore soaked kin larvae in a blend of synthetic SELF-1 peptides with the sequences identified from diverse conspecifics. These strains are recognized as unfamiliar and killed by the PS312 strain (*23*). These synthetic blends consisted of either the hypervariable regions (SELF-1 short peptide) or the hypervariable regions together with the preceding conserved region of SELF-1 (SELF-1 long peptide) from five genetically distant conspecifics. Incubation of PS312 animals in any of these unfamiliar SELF-1 peptide blends failed to disrupt kin recognition (Fig. 1E). It is possible that these synthesized peptides cannot be maintained in close proximity to the nematode surface, or alternatively, that SELF-1 may require additional post-translational modifications for activity.

We next sought to explore mechanisms of disrupting the surface itself. The nematode cuticle is an apical extracellular matrix that forms an external barrier to the environment. It is composed primarily of secreted collagens and cuticulins that assemble to form complex, three-dimensional structures (Fig. S2), with an overlying epicuticle and surface coat containing a variety of other non-collagenous proteins, lipids, and glycosylated molecules (*36–38*). To investigate the possibility that collagens or other cuticle-associated proteins are involved in kin signaling, we exposed *P. pacificus* larvae to either collagenases or a proteinase and conducted corpse assays. Exposing larvae to either one of these surface degradation methods for extended periods resulted in high levels of lethality. However, after non-lethal exposure times we observed no disruption to kin signaling abilities (Fig 1E). Thus, these components may be dispensable for the generation of the kin cues but we cannot exclude the possibility that the shorter, non-lethal exposure time used in our experiments was insufficient to degrade any putative signals. As lectins have previously been shown to bind surface carbohydrates in several nematode species and disrupt the exterior barrier function(*39–41*), we next assessed if they could disrupt kin-recognition signals in *P. pacificus*. Despite selecting lectins from numerous sources, we found that these also had no effect on kin-recognition, which was maintained at wildtype levels (Fig. 1E). To investigate the involvement of lipids or other amphiphilic molecules in kin signaling, larvae were soaked in either ethanol or the lipid-destabilizing cationic surfactant, cetyltrimethylammonium bromide (CTAB). Although lipids are mildly soluble in ethanol, treatment with low concentrations did not affect kin signaling (Fig. 1E). This may reflect experimental limitations, as washing in even relatively low concentrations of ethanol can be toxic and lethal to the animals. In contrast, washing with low concentrations of the surfactant CTAB significantly disrupted kin signaling (Fig. 1E). These results suggest that surface-associated lipids are essential for kin detection. Taken together, our findings indicate that the kin signal in *P. pacificus* is generally robust and resistant to various environmental challenges, but can be disrupted by agents that destabilize surface lipids.

### *self-1* mutants acquire aberrant surface lipids

It was recently shown that the environmentally exposed cuticular surface of both *C. elegans* and *P. pacificus* is dominated by an abundance of diverse lipids that differ between developmental stage and also show species-specific profiles (*28*). A similar finding was also observed in a parasitic nematode (*42*) . As *self-1* is so far the only identified kin-recognition signaling component, we next investigated if *self-1* mutants acquired a normal surface lipid profile using the same methodology. This required employing surface sensitive mass spectrometry, specifically the OrbiSIMS instrument, which combines a gas cluster ion beam (GCIB, Ar_3000_^+^) with an Orbitrap analyzer (*43*). We restricted our OrbiSIMS analysis to the outermost ∼50 nm of the *P. pacificus* cuticle, a depth that likely encompasses the surface coat and the epicuticle rather than the underlying collagen-rich cuticular layers (*36*) . This was achieved by following the protocol we developed recently (*28*) . Moreover, because the surface coat is thought to be compositionally dynamic and partially exchangeable with the environment (*44*), the detected signals likely represent a combination of epicuticular components and surface-associated secretions. (Fig. 2A).

**Figure 2.**
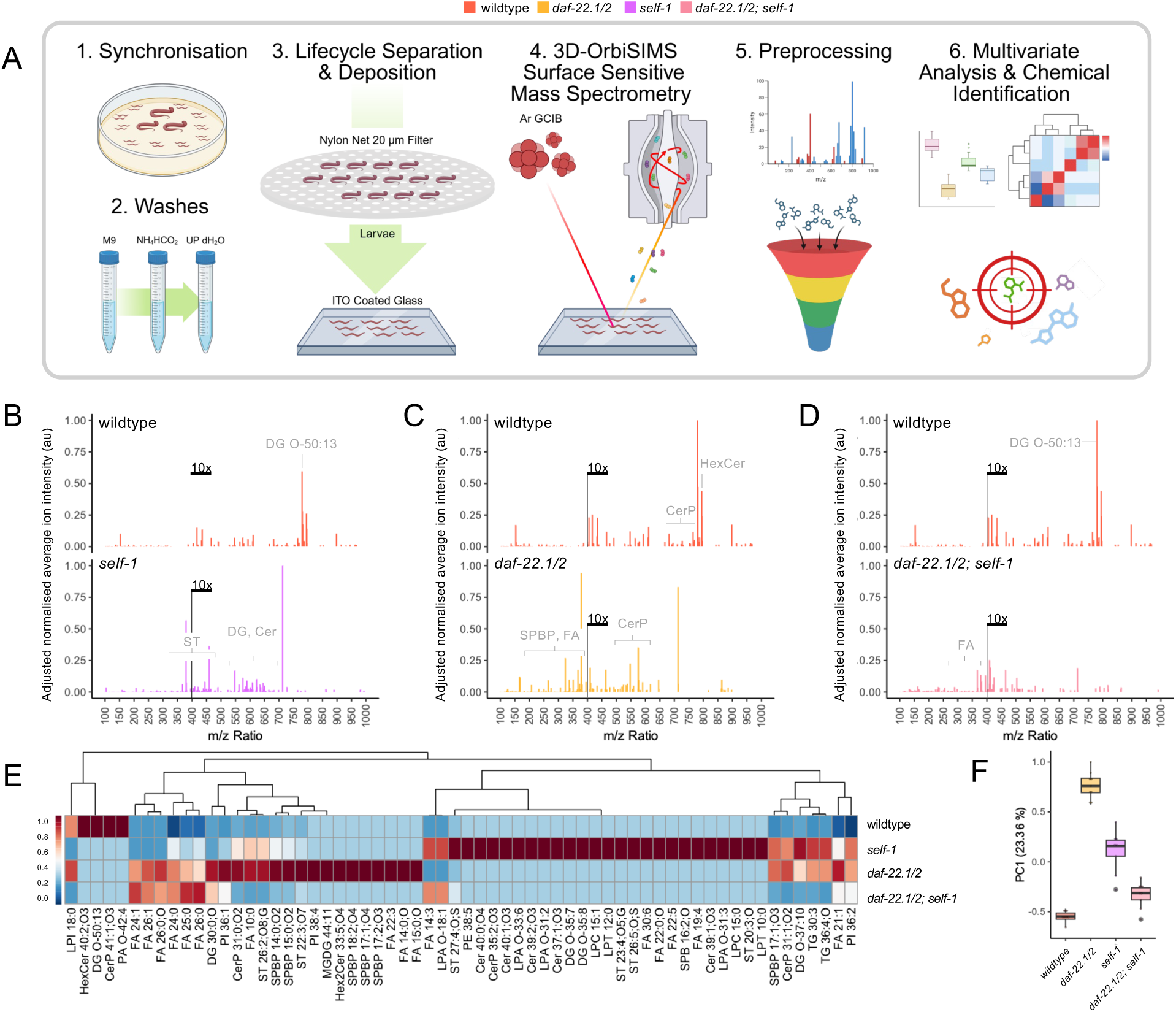
*self-1, daf-22.1/2* and *self-1;daf-22.1/2* mutants influence surface lipids through distinct pathways. (A) 3D OrbiSIMS workflow schematic. (B) Averaged surface secondary ion mass spectra exclusive to wildtype and *self-1,* (C) *daf-22.1/2* and (D) *self-1;daf-22.1/2* triple mutants *p* < 0.001 (***). (E) Hierarchical clustering heatmaps of distribution of exclusive lipids that were identifiable in lipid maps, for wildtype, and *self-1*, *daf-22.1/2*, and *self-1;daf-22.1/2* mutant animals, where phylogeny indicates shared presence or fragmentation of lipids and their relative presence on nematode surfaces. (F) Univariate PC1 wildtype vs mutant scores plot for wildtype and mutant larvae.

We focused on the larval stage as these animals are used as prey in our behavioral assays. We compared the averaged and normalized secondary ion spectra between wildtype and *self-1* mutants after eliminating background chemistries (Fig. S3) and analyzed the exclusive chemistries found on each. Strikingly, we identified a distinct surface chemical composition in *self-1* mutants. We observed a prominent absence of secondary ions between *m/z* 775-800 in *self-1* mutants, a region typically rich in triglycerides and diglycerides, such as DG O-50:13 (M-H, C_53_H_79_O_4_^-^, *m/z* 779.5976) in wildtype animals (Fig. 2B).

Additionally, *self-1* mutants exhibited an increase of mass ions ranging from *m/z* 350 to 750, consisting of diverse chemistries including sulphated sterols (e.g., ST 27:4;O;S), diglycerides (e.g., DG O-35:7, and DG O-35:8), and ceramides (e.g., Cer 39:1;O3, Cer 40:1;O3 and Cer 40:1;O4) (Fig. 2B and S3, S4).

Thus, *self-1* mutants present an atypical surface lipid composition and are kin-recognition defective.

In *C. elegans* and *P. pacificus, daf-22* mutants have been shown to acquire differential surface chemistries when compared to wild-type animals (*28*). In both species, *daf-22* is required to catalyze the terminal step in the peroxisomal β-oxidation pathway and is involved in the degradation of very long-chain fatty acids (*45–47*). In *P. pacificus* a recent gene expansion has resulted in two copies of *daf-22*, with one copy (*daf-22.1*) contributing the majority of the catalytic function, while the second copy (*daf-22.2*) has a reduced role (*47*). A cumulative effect is observed in double mutants, hereafter referred to as *daf-22.1/2*. In *P. pacificus, daf-22.1/2* mutants have a distinct surface profile from both wildtype animals and *self-1* (Fig 2C). This includes the presence of different fatty acids (e.g., FA 22:3, FA 14:0;O and FA 15:0;O), sphingoid base phosphates (e.g., SPBP 18:2;04, SPBP 17:1;04 and SPBP 17:2;03) and ceramide phosphates (e.g., CerP 31:0;O2) (*28*). While there are differentiating lipids between *self-1* and *daf-22.1/2*, they also share similarities, including ceramide phosphates (CerP 31:1;O3) and triglycerides, (e.g. TG 30:3 and TG 36:4;O). Furthermore, by generating a *self-1; daf-22.1/2* triple mutant we uncovered a combination of unique and cumulative effects of these mutations resulting in discrete surface chemical compositions (Fig. 2D and 2E). These findings were further supported by hierarchical clustering (Fig. 2E) and PCA analysis (Fig. 2F and S4). Notably, in *self-1; daf-22.1/2* triple mutants, we observed a marked absence of many complex chemistries compared to wild type as well as *self-1*, and *daf-22.1/2* mutants (Fig. 2E). In addition, all mutant strains exhibited novel chemistries absent in wildtype animals and all mutants were absent of CerP 41:1;O3, phosphatic acid PA O-42:4, hexosylceramide HexCer 40:2;O3 and diglyceride DG O-50:13 (Fig. 2B-E). Together, these data indicate that *self-1* and *daf-22* are both required for the surface lipid profile in *P. pacificus*, with each mutant displaying overlapping but distinct surface features.

### Abnormal surface lipids correlate with kin-recognition defects

Given the aberrant surface profiles observed in both *P. pacificus self-1* and *daf-22* mutants, we next assessed if *daf-22* mutants are also kin-signaling defective. We conducted corpse assays on both *daf-22.1/2* and the *self-1;daf-22.1/2* triple mutants and compared this to wildtype and kin-recognition defective *self-1* mutants (*23*). Wildtype predators failed to detect *daf-22.1/2* mutants as kin and instead these were killed at a similar frequency to *self-1* mutants (Fig. 3A). Furthermore, *self-1;daf-22.1/2* triple mutants were also kin-signaling defective, although this was not significantly different from either the *self-1* or *daf-22.1/2* single mutants under these assay conditions (Fig. 3A). To further investigate the disruption of kin-recognition in these mutants, we analyzed the behavioral states of predators while surrounded by mutant prey. We previously developed an automated behavioral tracking and machine learning model to identify behavioral states associated with feeding and predatory activity (*48*, *49*). This approach enabled us to identify six behavioral states associated with the *P. pacificus* behavioral repertoire (*49*) . Three of these are similar to behavioral states observed in *C. elegans* including a ‘dwelling’ state and two ‘roaming’ states (*50*). The remaining three are predation-associated states and include ‘predatory search’, ‘predatory biting’, and ‘predatory feeding’ (Fig. 3B). Importantly, the two most predictive behavioral features of predation states are velocity and pumping rate, with predatory biting and feeding occurring at near-zero velocities. By observing the joint distribution of these two features, it is possible to visualize the prevalence of the predatory and non-predatory states (Fig. 3B). Additionally, we can determine behavioral state occupancy while animals are surrounded by potential prey using a machine learning model previously trained on behavioral data using both unsupervised and supervised methods (Fig. 3B)(*49*). We employed these methods to our mutants and we observed no predatory behavioral states when predators were surrounded by kin, but robust predatory state occupancy while predators were in the presence of *C. elegans* larvae (Fig. 3C-D). Crucially, when predators were exposed to larvae of either *self-1*, or *daf-22.1/2* single mutants or *self-1;daf-22.1/2* triple mutants, we observed an increase in ‘predatory search’ state occupancy indicating an increased environmental sampling (Fig. 3D). Furthermore, statistically significant occupancy of the ‘predatory biting’ state was detected when predators were surrounded by *self-1;daf-22.1/2* triple mutants (Fig. 3D). This was accompanied by increased transitions into these behavioral states (Fig. 3E). Thus, our results reveal *daf-22* mutants are kin-recognition defective and that *P. pacificus* predators enter more aggressive predatory states when surrounded by animals with a weakened kin-recognition signal.

**Figure 3.**
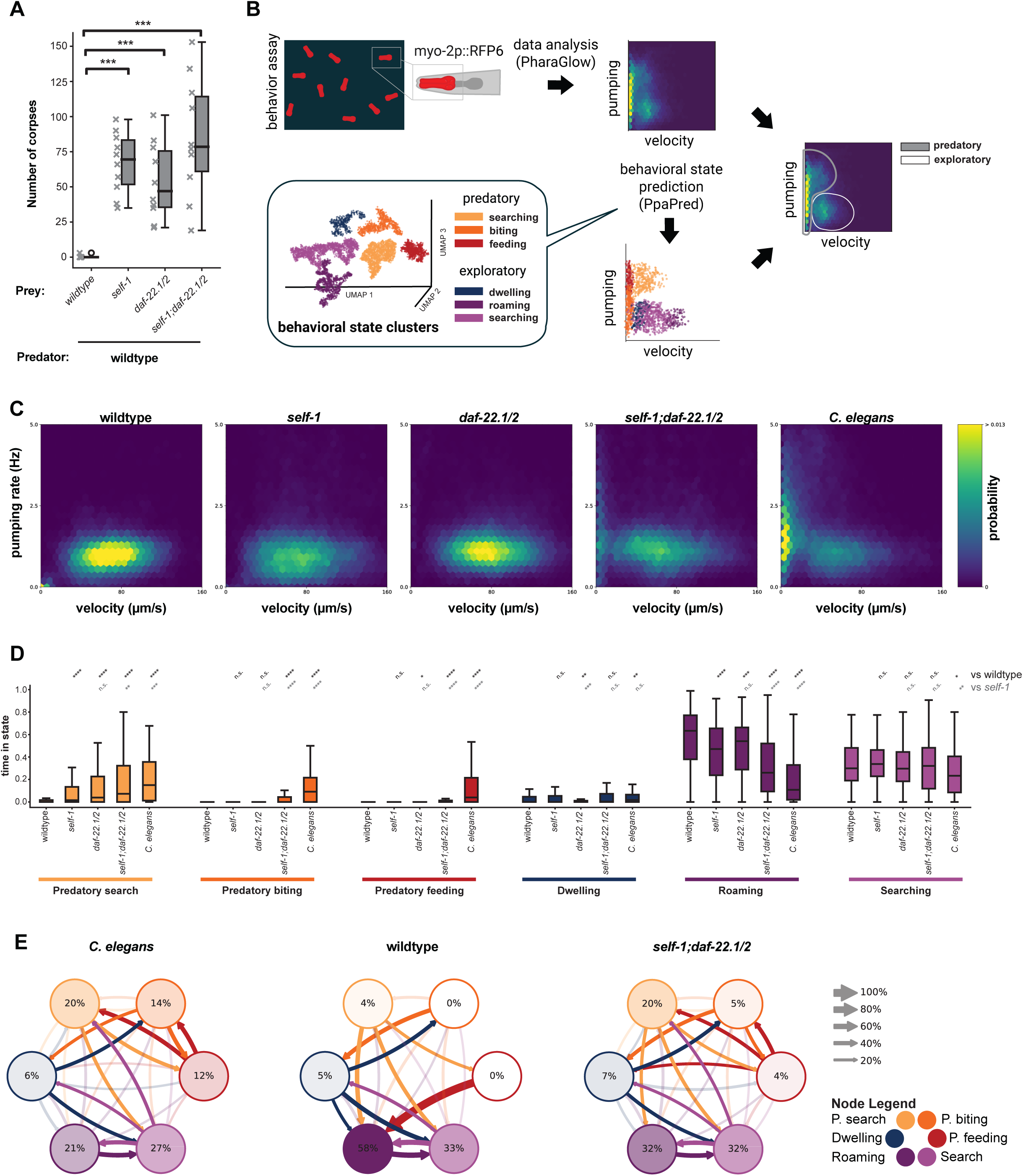
Abnormal surface lipids are associated with kin-recognition defects. (A) Number of corpses present after 24 h when wildtype predators are exposed to mutant prey. (B) Schematic of automated behavioral tracking and machine learning model to identify behavioral states. (C) Probability density plot of wildtype velocity and pumping rate when exposed to corresponding prey (identified above each plot). (D) Relative time spent in each behavioral state normalized to total track duration. Significance assessed using Mann-Whitney-U-Test with Bonferroni correction for multiple comparisons. Statistical tests shown in black correspond to data compared with WT control (PS312), while those shown in grey correspond to data compared with *self-1*. (E) Average transition rates between behavioral states for WT *P. pacificus* predators on *C. elegans*, *P. pacificus*, and *self-1;daf-22.1/2* mutant prey.

## Discussion

Molecular determinants of kin-recognition have remained elusive in many multicellular organisms. However, in the nematode *P. pacificus,* genetic components regulating these vital systems are beginning to be uncovered. In *P. pacificus,* kin-recognition has evolved alongside predation as a mechanism to avoid the costly killing of their own progeny and close relatives (*22*, *23*). These behaviors are contact mediated, with prey identity determined through surface-associated interactions (*33*, *34*). While these signals depend on a small peptide encoded by *self-1,* the precise signaling mechanism is currently unknown (*22*, *23*). The outer 50 nm of the nematode cuticle is predominantly composed of lipids (*28*), and we found that disrupting this chemical group by a surfactant wash induced kin-recognition defects in *P. pacificus*. In addition, mutants with abnormalities in surface lipid chemistries were also unable to present a robust kin-recognition signal. These issues may indicate that the surface lipids themselves act as signaling components. Alternatively, the kin signal could be either secreted into the lipid layer or be embedded through lipid binding, and thus maintained in close proximity to the organism’s surface. Furthermore, these mechanisms are not mutually exclusive and a combination of these factors may work together to act as the kin cue.

As surface lipids in nematodes create a hydrophobic barrier preventing the organism’s desiccation and protecting it from other external environmental damage (*51*, *52*), an alternative hypothesis is that significant deterioration in the lipid layer may result in cuticular or other structural damage, inducing the observed kin-recognition defect. Importantly, the surface properties of *P. pacificus* may serve functions analogous to the waxy cuticular hydrocarbon (CHC) layer that surrounds many insect species. These CHCs prevent the insect’s desiccation and can also communicate information including the sex, age and reproductive status as well as nestmate identity as a surrogate for kinship (*53*). The formation of insect CHCs also relies on well-characterized fatty acid biosynthesis pathways (*18*), and are detected by specialized odorant receptors (*54*). Our findings suggest a similar mechanism may exist in nematodes, where the peroxisomal β-oxidation pathway, responsible for degrading very long chain fatty acids, may also play an important role in establishing surface signals. Taken together, these data support the notion that lipids play a significant role in enabling kin-identity across several taxa.

Finally, we observed an unexpected disruption of the surface-lipid profile in *self-1* mutants. *self-1* encodes a small peptide which consists of distinct domains including a hypervariable domain at the C-terminus that has been shown to be essential for its function. This peptide shows striking diversity between strains and thus, is thought to contribute to strain-specific kin signals (*23*). However, some strains with identical hypervariable regions still cannibalize one another indicating a more complex mechanism than first predicted (*22*). These data indicate that surface lipids are an additional element necessary for establishing the kin signal and importantly, *self-1* is necessary to generate a normal surface lipid profile. Despite this, we note that predation levels on lipid-defective conspecifics, including *self-1* and *daf-22* mutants, still do not approach the levels of predation observed when exposed to another distantly related species such as *C. elegans* (*30*) . This suggests that additional components act together to contribute to generating an effective kin signal.

Therefore, given the contact-mediated nature of kin signaling and the highly complex chemical composition of the outermost cuticle, we predict that the putative kin signal likely involves multiple other cuticular elements in addition to lipids and it is tempting to speculate that the cuticle structure itself may function as part of the kin-signaling system. Consistent with this, recent studies in *C. elegans* have demonstrated that the mechanical properties of its surface are important for contact-dependent cues including mate recognition (*55*). Thus, our work here demonstrates that in predatory nematodes the external surface composition is associated with sophisticated chemical cues, including for establishing kin-identity, and reveals the nematode surface as a signaling organ regulating contact-dependent behaviors.

## Materials and Methods

### Behavioural methods

#### Nematode culture

All nematodes used were maintained on standard NGM plates on a diet of *Escherichia coli* OP50.

##### Nematode strains

| Name | Strain | Alleles | Species |
| --- | --- | --- | --- |
| N2 | N2 | wildtype | *C. elegans* |
| PS312 | PS312 | wildtype | *P. pacificus* |
| PS1843 | PS1843 | wildtype | *P. pacificus* |
| *self-1* | RS3246 | *self-1 (tu1112)* | *P. pacificus* |
| *daf-22.1; daf-22.2* | RS2770 | *daf-22.1 (tu489); daf-22.2 (tu504)* | *P. pacificus* |
| *self-1; daf-22.1; daf-22.2* | JWL90 | *self-1 (bnn78); daf-22.1 (tu489); daf-22.2 (tu504)* | *P. pacificus* |

#### Corpse assays

We used corpse assays to rapidly quantify predatory behaviors between different strains. Prey were maintained on NGM plates seeded with OP50 bacterial food until freshly starved, resulting in an abundance of young larvae. In order to isolate a pure culture of larvae, the plates were washed with M9 and filtered through 20 µm mesh into a tube. The tube is centrifuged in order to get a pellet of larvae, and either 1.5 µl of *P. pacificus* or 1.0 µl of *C. elegans* prey is pipetted onto an unseeded 6 cm NGM plate.

Predatory young adult nematodes are screened for appropriate mouth form and five are added on to each assay plate. After 24 h for *P. pacificus* prey and 2 h for *C. elegans* prey, the plates were subsequently screened for corpses.

#### Environmental stressor treatments

Treatment conditions:

Starvation: prey is collected and starved on an unseeded plate until used for assays.

Treatments: all conditions were treated on a rotator at room temperature unless mentioned otherwise. After each treatment, the worms were washed once with M9, collected by centrifugation and pipetted onto unseeded NGM plates for corpse assays.

NaCl: NaCl was diluted with M9 to 200 mM, 300 mM and 400 mM, and worms were treated for 1 h in each concentration.

Heavy metals: Cu^2+^: 1 mM CuCl_2_ was diluted with M9. Zn^2+^: 3 mM ZnCl_2_ was diluted with M9. Prey was treated with the solution for 3 h.

Collagenases: Collagenase IV was prepared with 10 mg/ml concentration in M9, and prey was treated with the solution for 1 h. Collagenase VII was prepared with 5 mg/ml in M9. Prey was treated with the solution for 30 min at 30°C.

Cysteine proteinase: prey was treated with 50 µM in M9 for 3 h.

Short/long peptides: blend of SELF-1 peptides from conspecifics containing hypervariable regions only (short) or hypervariable region together with the preceding conserved region of SELF-1 (long) were diluted to 10% with M9 with final volume of 100 µl, and prey was treated for 1 h. Each of the peptide blends consist of SELF-1 peptides from following strains:

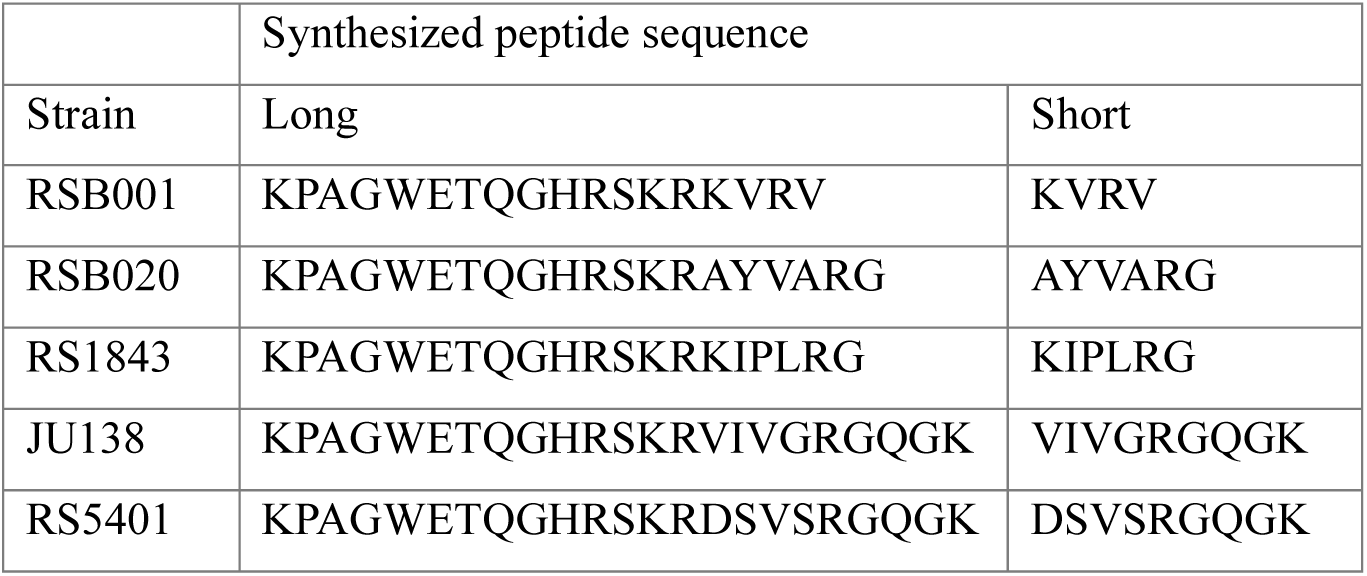

Lectins: each lectin was prepared as 1mg/ml concentration in M9, the prey was treated in the solution for 1 h.

EtOH: prey was treated with 10% EtOH diluted in M9 for 30 min. CTAB: prey was treated with 0.25% CTAB diluted in M9 for 15 min.

As several treatments resulted in a large number of dead animals when exposure times were extended, treatment times were dependent upon and modified according to the overall health of the assayed prey.

### CRISPR/Cas9 induced mutations

CRISPR/Cas9 was used to induce mutations in candidate genes with gene specific crRNA and universal trans-activating CRISPR RNA (tracrRNA) purchased from Integrated DNA Technologies (IDT). 5 µl of each 100 µM stock were mixed and denatured at 95 °C for 5 min, then cooled at room temperature to anneal. Cas9 (IDT) was added to the hybridized product and incubated at room temperature for 5 more min. The mixture was diluted with TE buffer to a final concentration of 18.1 µM sgRNA and 2.5 µM Cas9. This was injected into the germlines of the required *P. pacificus* strains. Eggs from injected P0s were recovered up to 16 h post injection. Once these eggs have hatched and the worms were old enough to be picked, the F1s were segregated onto individual plates until they have developed and laid eggs. F1 animal genotypes were analyzed via Sanger sequencing to identify mutations, and those worms with mutations were further re-isolated in homozygosis.

### Automatic behavioral tracking and machine learning

Behavioral tracking and behavioral state predictions for the predators were performed as previously described (*49*). Briefly, prey to be assayed were collected as described under corpse assay, and 3 µl of the prey pellet was pipetted onto the assay plate. Once the prey has spread across the plate, a Copper arena (1.5 cm x 1.5 cm) was placed in the center of the assay plate. 40 adult worms with an integrated *myo-2p*::RFP were starved for 2 h prior to the experiment on an NGM plate and were subsequently added to the assay arena. The animals were left to acclimate to the environment before being recorded at 30 fps for 10 min. An epi-fluorescence microscope (Axio Zoom V16; Zeiss) with a Basler camera (acA3088-57um; BASLER) attached was used for recording. Animals were then tracked using python analysis package PharaGlow (*48*) adapted for *P. pacificus*. To acquire locomotion velocity and pumping rate, post-processing was conducted with a previously developed python script. Furthermore, python package PpaPred was used to automatically predict the behavioral states of the predators.

### Statistical analyses

The statistical analysis of the corpse assays was performed using Mann-Whitney-U test, with wildtype condition as control and Bonferroni correction was applied. P-value < 0.05 (*), p-value < 0.01 (**), p-value < 0.001 (***), p-value < 0.0001 (****), and pairs without stars are not significant.

For the statistical analysis of the state predictions (relative time in state, mean bout duration, transition rates), a two-tailed Mann-Whitney-U test was used. The statistical analysis was performed on each condition and state to its respective control. A Bonferroni correction was applied to take into account that the probability of a false positive significant test rises with the number of compared conditions.

Therefore, the *p* values reported are corrected for the number of comparisons made to the respective control. All raw *p* values and sample sizes are available in Table S1. The boxplots follow Tukey’s rule where the middle line indicates the median, the box denotes the first and third quartiles, and the whiskers show the 1.5 interquartile range above and below the box. In all figures p-value ≥ 0.05 (n.s.), p-value < 0.05 (*), p-value < 0.01 (**), p-value < 0.001 (***), and p-value < 0.0001 (****).

### Surface chemistry methods

#### OrbiSIMS

Calibration of the Orbitrap analyzer was performed using silver clusters, according to the protocol described by Passarelli et al. The Bi ^+^ liquid metal ion gun and ThermoFisher Tune software were executed for calibration. An Ar ^+^ primary gas cluster ion beam (GCIB, 20 keV, 2 μm diameter, duty cycle set to 27.78%, and target current was 24 pA) was used for sample sputtering and ejection of secondary ions. The Q Exactive images were acquired using a random raster mode (field of view 300 × 300 μm, pixel size 5 μm, cycle time 200 μs, and optimal analyzer target −69.5 V). Argon gas flooding was in operation to facilitate charge compensation regulating the pressure in the main sample chamber to 9.0 × 10^−7^ bar throughout the analysis. The images were collected in negative polarity (*m*/*z* 75–1125) with constant injection time (500 ms), total ion dose per measurement (2.70 × 10^14^ ions/cm^2^), and mass-resolving power (240,000 at *m*/*z* 200). Given the sputtering rate for organic materials, an ion dose of 3.00 × 10^14^ ions/cm^2^ was estimated to have analyzed a sample depth of approximately 50 nm.

#### Comparative Peak Analyses

Student’s *t*-test (two-tailed, unequal variance) was performed to identify significantly different mass during binary comparisons of data sets. Standard deviation was calculated between ROI (*n* = 9) and plotted as error on accompanying bar charts. Significantly different data were categorized based on their *p*-value thresholds such that p-value < 0.001 (***). The following lipid abbreviations are used: FA, fatty acid; DG, diacylglycerol; TG, triacylglycerol; Cer, ceramide; CerP, ceramide phosphate; HexCer, hexosylceramide; Hex2Cer, dihexosylceramide; SPBP, sphingoid base phosphate; ST, sterol sulphate; PA, phosphatidic acid; PE, phosphatidylethanolamine; PI, phosphatidylinositol; LPA, lysophosphatidic acid; LPI, lysophosphatidylinositol; LPC, lysophosphatidylcholine; LPT, lysophospholipid; MGDG, monogalactosyldiacylglycerol.

#### Principal Component Analyses

Principal Component Analysis (PCA) was conducted on comparative data sets, scaling, standardizing, and centering each variable so that they have a standard deviation of 1 and a mean of 0, respectively. This step was important to ensure that all variables contribute equally to the principal components, eliminating any undue influence from variables. For 2D and 3D scores, plots ellipses were used to visualize the 95% confidence interval to provide a guide to the consistency of data groupings.

#### Hierarchical Clustering Heatmap Analyses

Hierarchical clustering heatmap analysis utilized a data reduction approach by selecting only mass ions with principal component loadings that were greater than one standard deviation from the mean. This data reduction was key to data visualization, ensuring a focused analysis on the organisms and *m*/*z*-based separation of variables. Visualizations were optimized though the application of heatmaps for columns of data comparing strains.

#### Data Analysis and Packages

A range of R packages were used, each serving a specific purpose. SurfaceLab (Version 7) was used for data acquisition and manipulation, while R Studio (2022.07.02 + 576) provided an integrated development environment for R, the programming language version 4.2.2. The “dendextend” package (version 1.17.1) aided in visualizing and comparing hierarchical clustering trees. Data frame tools were handled by dplyr (version 1.1.2), and factoextra (version 1.0.7) and factomineR (version 2.8) were used for visualization and extraction in multivariate analysis. The study also employed “forcats”s, “ggforce” (version 0.4.1) to accelerate “ggplot2” (version 3.4.3), and “ggrepel” (version 0.9.3) for positioning nonoverlapping labels in “ggplot2”. Additional packages included “ggsignif” (version 0.6.4), “ggtext” (version 0.1.2), and “gplots” (version 3.1.3) for plotting enhancements, “heatmaply” (version 1.4.2) for interactive cluster heat maps, and “lattice” (version 0.20–45) for trellis graphics. The “pheatmap” package (version 1.0.12) was used for heatmap production, and “plotly” (version 4.10.2) was used for interactive web graphics. The study also utilized “readr” (version 2.1.4) for reading rectangular data, “scales” (version 1.2.1) for scaling and formatting axis labels, “scatterplot3d” (version 0.3–44) for 3D scatter plots, “stringr” (version 1.5.0) for string operations, “svglite” (version 2.1.1) for SVG graphics, “tibble” (version 3.2.1) as a modern data.frame variant, and “tidyr” (version 1.3.0) for data tidying. The “tidyverse” collection (version 2.0.0) provided a comprehensive suite of data science packages, while “viridis” (version 0.6.3) and “viridisLite” (version 0.4.2) were used for creating perceptually uniform color maps.

## Acknowledgements

We would like to thank all the Lightfoot lab, Prof. Morgan R. Alexander and Dr. David J. Scurr for useful discussions as well as Monika Scholz for critical reading of the manuscript. We thank the Sommer lab (MPI for Biology, Tübingen) for providing lab strains PS312 and RS2770.

## Author contributions

JWL and VC developed the concept and design of the project, and secured funding. FH, DLG, AMK, NZ and VC assisted in experimental design, performed research, and collected data. Surface chemistry data analysis was performed by VC. Behavioural data analysis was performed by FH and DLG. FH and VC compiled data and designed figures. FH, DLG, VC and JWL drafted the original version of the manuscript. FH, DLG, AMK, VC, and JWL and edited and revised the manuscript.

