## Supplemental Files for "Contact-based kin discrimination is associated with specific surface lipids in the cannibalistic nematode *Pristionchus pacificus*"

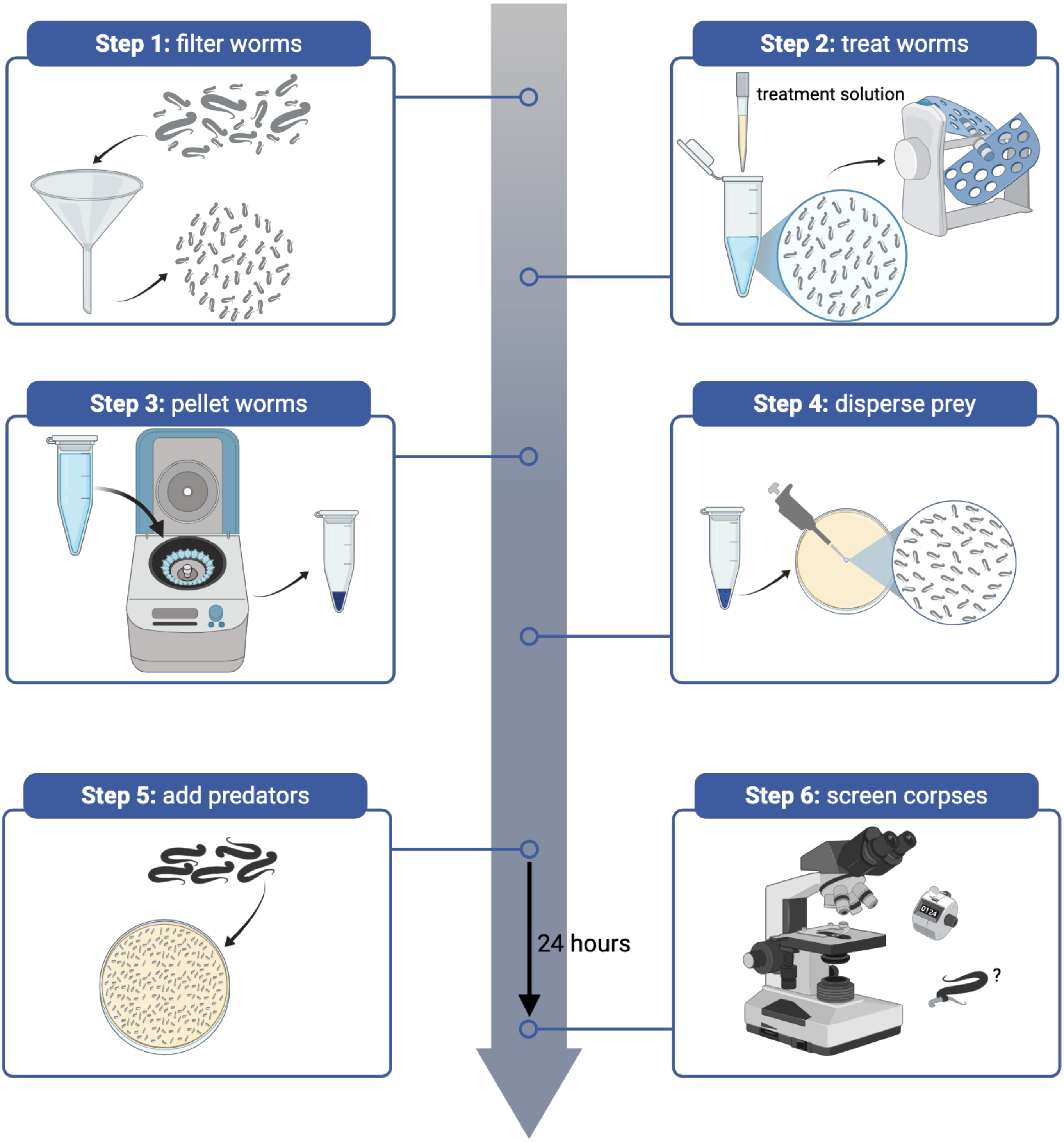


**Figure S1. Schematic of environmental stressor treatment and corpse assay method.** When corpse assays were performed without treatment, step 2 is omitted from the procedure.

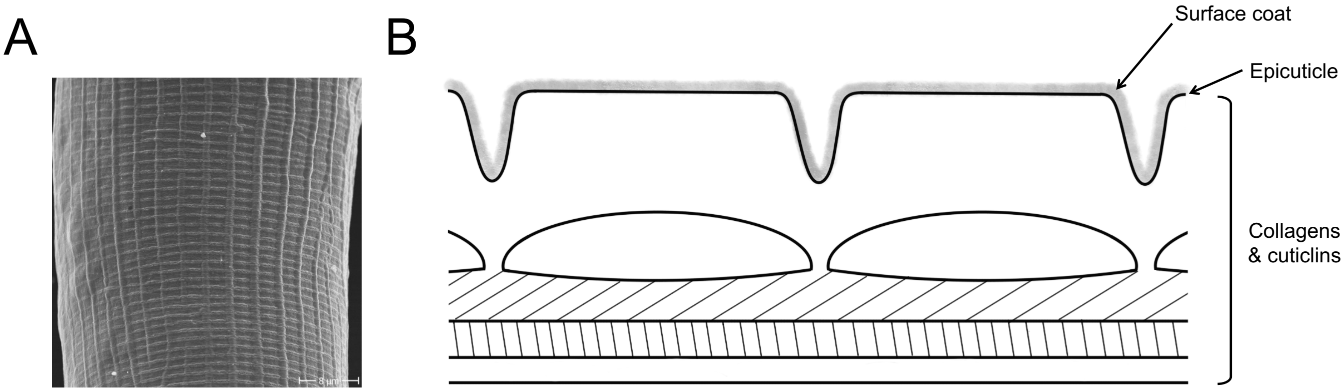


**Figure S2. *Pristionchus pacificus* surface structure.** (A) Scanning electron micrograph of *P. pacificus* cuticle. (B) Schematic depicting the structural features of the cuticle as seen in a longitudinal section. Superficially, the thin surface coat and epicuticle line a multi-layered, apical extracellular matrix composed primarily of collagens.

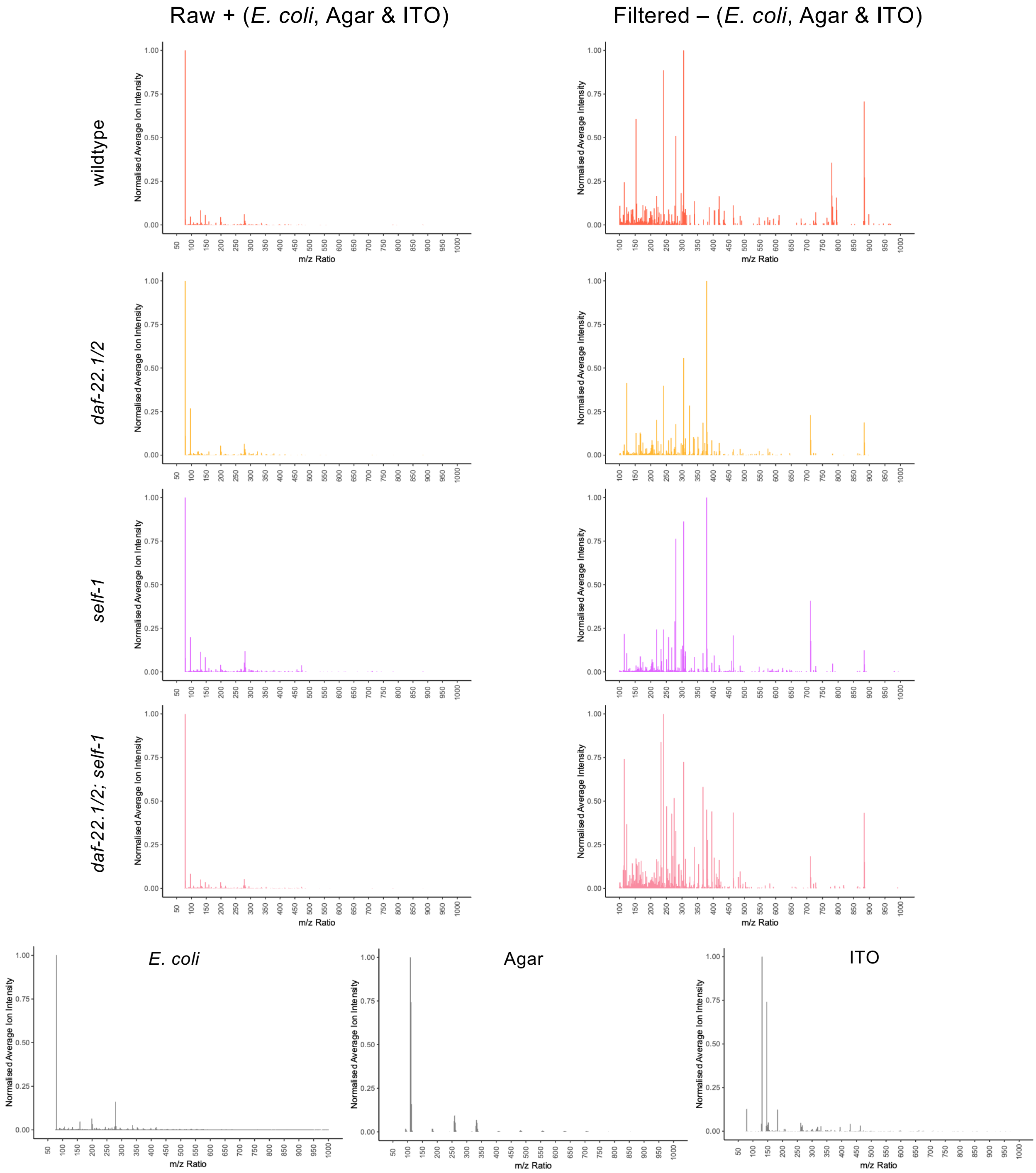


**Fig. S3 | *P. pacificus* mass spectra optimisation.** Raw and filtered secondary ion mass spectra, eliminating *E.coli*, Agar and ITO, ions to 5 ppm.

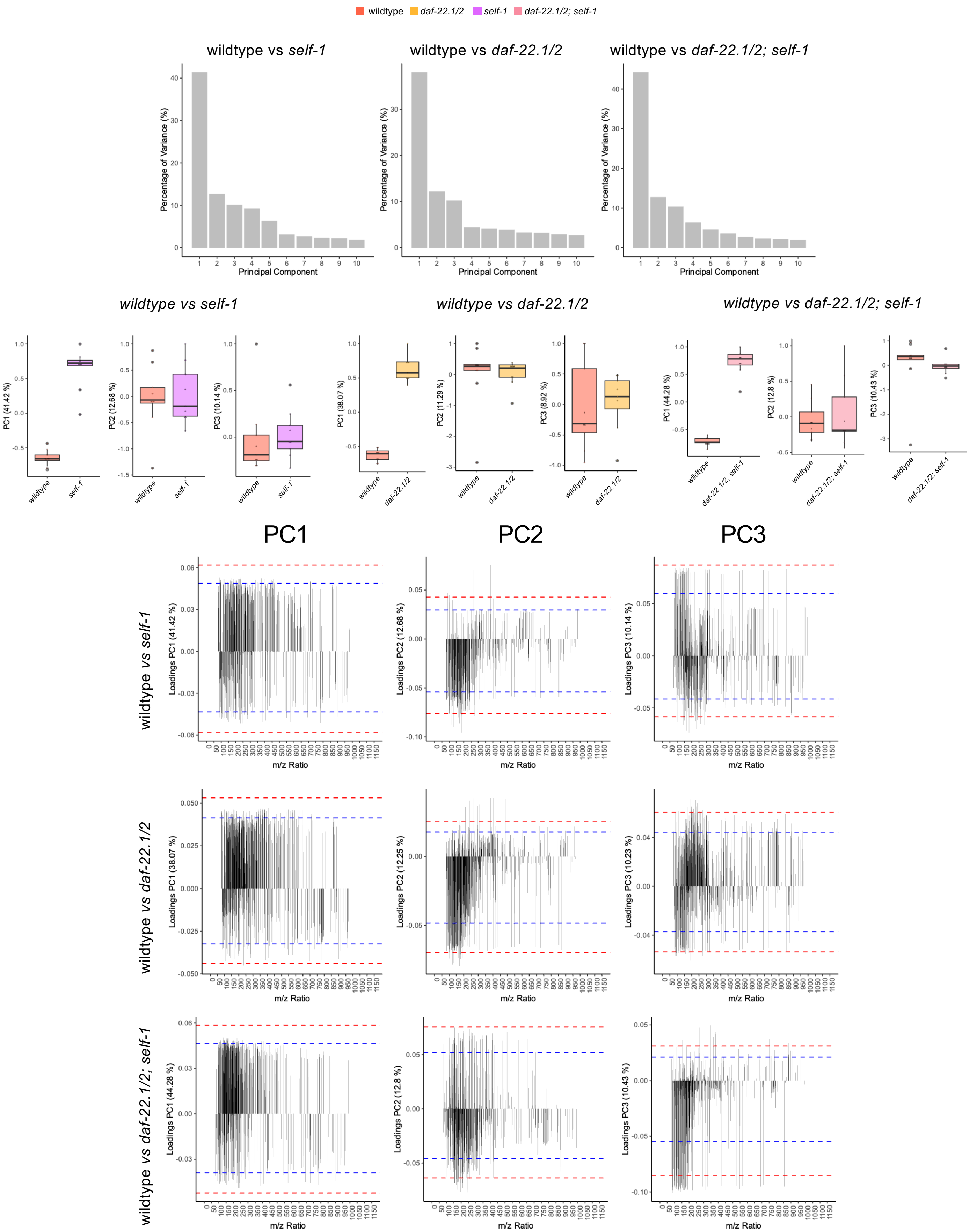


**Fig. S4 | Binary PCA analysis of *P. pacificus* wildtype vs *mutants daf-22.1/2, self-1, daf-22.1/2; self-1*. a** Percentage of variance on each principal component, b Univariate PCA 1-3 scores plot and corresponding loadings where blue and red dashed lines indicate 1 and 2 standard deviations from the mean, respectively.

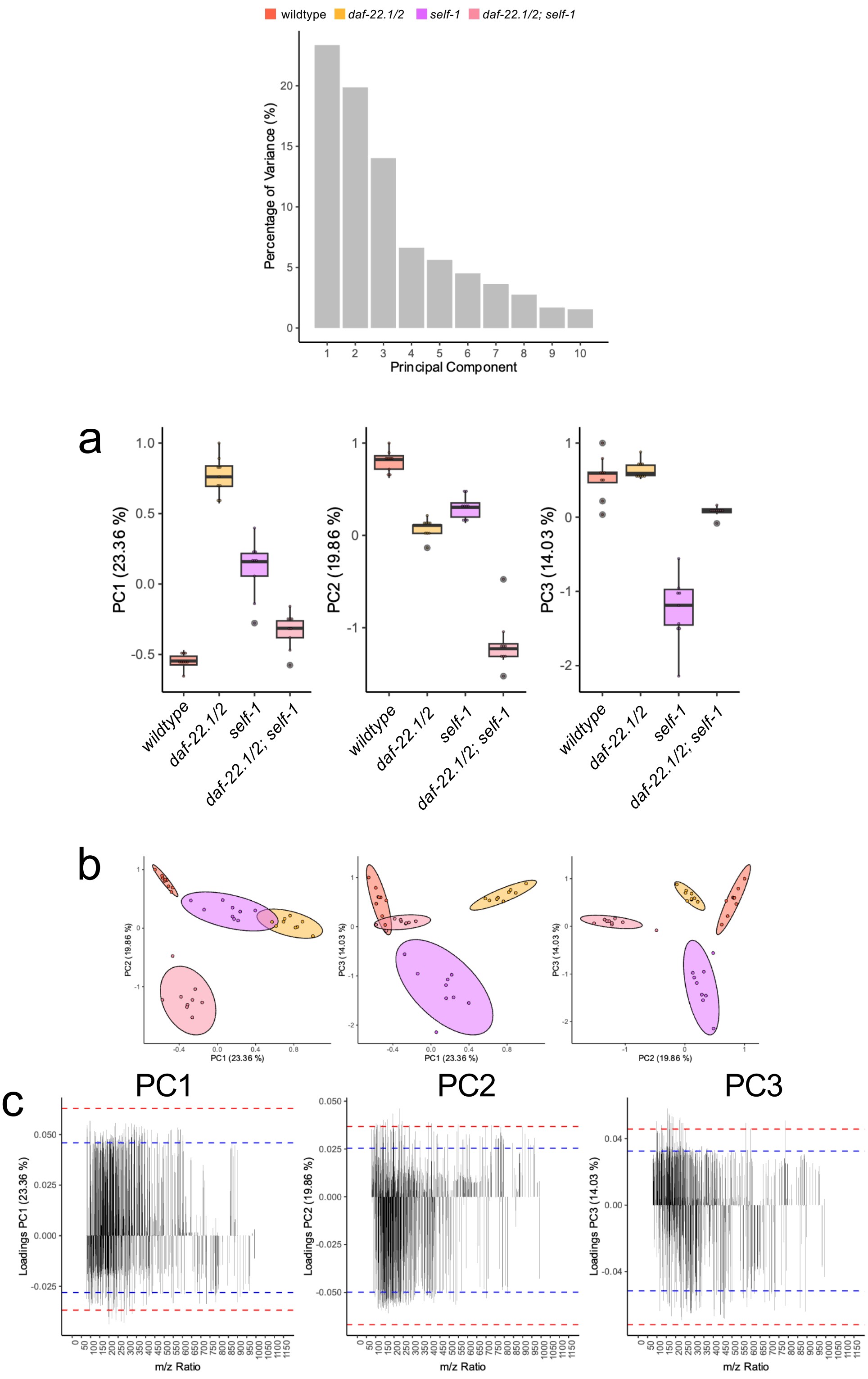


**Fig. S5 | Multivariate analysis of *P. pacificus* wildtype vs *mutants daf-22.1/2, self-1, daf-22.1/2; self-1*. a** Percentage of variance on each principal component. **b** Univariate and **c** multivariate PC 1-3 scores plot. d PC 1-3 loadings plots, where blue and red dashed lines indicate 1 and 2 standard deviations from the mean, respectively.
